# PI-FC: Pre-training Individual-specific Functional Connectome through State-invariant Contrastive Learning

**DOI:** 10.1101/2025.09.21.677570

**Authors:** Yingjie Peng, Xiaohan Tian, Yini He, Shangzheng Huang, Tongyu Zhang, Junxing Xian, Tian Gao, Qi Wang, Changsheng Dong, Xiya Liu, Kaixin Li, Yao Ge, Xianchang Zhang, Lei Wang, Yiheng Tu, Bing Liu, Meiyun Wang, Yan Yan, Ang Li

**Affiliations:** State Key Laboratory of Cognitive Science and Mental Health, Institute of Biophysics, Chinese Academy of Sciences, Beijing 100101, China; University of Chinese Academy of Sciences, Beijing 100049, China; State Key Laboratory of Cognitive Neuroscience and Learning, Beijing Normal University; Beijing 100091, China; Department of Medical Imaging, Henan Provincial People’s Hospital & the People’s Hospital of Zhengzhou University, No. 7 Weiwu Road, Zhengzhou, Henan, 450003, China; MR Research Collaboration, Siemens Healthineers Ltd., Beijing, China; Shenzhen Institute of Advanced Technology, Chinese Academy of Sciences, Shenzhen, Guangdong, 518055, China; State Key Laboratory of Cognitive Science and Mental Health, Institute of Psychology, Chinese Academy of Sciences, Beijing 100101, China; Biomedical Research Institute, Henan Academy of Sciences, Zhengzhou, Henan, 450046, China

## Abstract

Functional MRI enables non-invasive mapping of brain connectivity, yet its clinical translation remains hindered by uncontrolled state-dependent variability that obscures individual-specific signatures during routine scanning. Here we introduce PI-FC — a deep learning framework leveraging state-invariant contrastive learning to extract stable individual brain signatures across diverse arousal levels, cognitive states, and temporal scales spanning tens of seconds to hours. PI-FC achieves equivalent phenotypic prediction accuracy using substantially reduced scanning time, and eliminates state-dependent effects varying task demands and brain states. Trained on 36,119 subjects across 8 independent datasets, our model demonstrates superior cross-site generalization and outperforms traditional functional connectome (FC) in predicting neuropsychiatric conditions including schizophrenia, autism, depression, and anxiety. Furthermore, PI-FC enables zero-shot inference of brain age, biological sex, and cognitive ability without site-specific retraining. Overall, PI-FC represents a robust, clinically scalable framework that overcomes fundamental barriers to real-world deployment of precision functional neuroimaging.

## Introduction

The advent of large-scale neuroimaging initiatives has transformed neuroscience by enabling brain-wide association studies (BWAS)^1–3^ and precision medicine approaches^4,5^. Among the most promising neuroimaging-based metrics is the functional connectome^6,7^, which quantifies inter-regional neural synchronization over time. FC-based analyses have proven valuable for characterizing individual differences across multiple domains, including biological sex^8^, developmental and aging trajectories^9^, cognitive abilities^10^, and various neuropsychiatric conditions such as autism spectrum disorder^11^, schizophrenia^12^, and major depression^13^. Recent work has further demonstrated that this capacity to capture individual-specific signatures^14^ generalizes across diverse brain states, extending FC analyses extended beyond resting-state to task-based and naturalistic paradigms^15,16^.

Despite its promise, FC estimation from clinically-feasible protocols faces significant barriers to clinical translation due to uncontrollable state-dependent fluctuations. Standard 8-15 minute resting-state paradigms exhibit substantial FC variability due to multiple temporal factors^17–19^: approximately 30% of subjects struggle to maintain stable wakefulness beyond 3 minutes^20–22^, while spontaneous cognitive activity induces hidden state transitions that destabilize FC measurements across sessions^23–25^. Although research-grade extended scanning protocols (> 30 minutes across sessions) can achieve robust individual-level FC estimation^18,26^ with superior phenotypic prediction accuracy^6,27^, such approaches incur considerable costs approaching US$1,000 per hour^28–30^, and preclude acquisition of other crucial neuroimaging modalities within feasible scanning sessions^31^. Task-based paradigms partially address state variability by constraining brain activity through specific task-imposed cognitive demand, but introduce standardization and generalization challenges across diverse populations such as patients, children, and elderly individuals^32–34^. Moreover, integrating data across different paradigms (e.g., resting-state, various tasks) for a unified analytical framework remains technically challenging^33,35^.

Emerging evidence suggests the existence of a stable, state-invariant FC architecture underlying various brain states^36,37^. Indeed, composite FC measures combining resting-state and task fMRI exhibit enhanced test-retest reliability and superior predictive utility compared to resting-state FC alone^19,27,38^. The individual-specific nature of FC is further evidenced by its capability to serve as a neural ‘fingerprint’ across different scanning sessions and paradigms^14,39^, albeit with higher identification accuracy within resting-state scans than between different task conditions^14^. Moreover, recent supervised approaches have achieved accurate individual identification using merely 20 seconds of resting-state data, suggesting the preservation of individual-specific FC information across brief timescales^40^. However, these methods rely heavily on multiple sessions from the same individual. In parallel, contrastive and self-supervised learning have been applied to FC networks derived from resting-state fMRI^41–43^, but these efforts have rarely attempted cross-dataset generalization and have not explored short-timescale FC enhancement. The critical challenge remains: generalizing to entirely new individuals and extracting reliable individual patterns from single brief acquisitions used in clinical practice^30,44,45^.

To address these challenges, we propose PI-FC (**P**re-trained **I**ndividual-specific **F**unctional **C**onnectome), a deep learning framework that mitigates major sources of FC variability—such as scan duration, arousal levels, meta-states, and task conditions—to extract stable individual signatures from multi-session fMRI data and generalize to single-scan acquisitions from independent subjects (**Fig. 1a**). Our core hypothesis employs state-invariant contrastive learning to transform FC representations from a common space dominated by state-dependent fluctuations into a new embedding space that preserves individual-specific patterns (**Fig. 1b**). To implement this transformation, the PI-FC framework (**Fig. 1c**) leverages a key insight: individual-specific connectivity patterns remain consistent across sessions while state-dependent fluctuations vary unpredictably across different scanning conditions. Through dense temporal sampling and contrastive learning, the framework learns to distinguish stable individual signatures from transient fluctuations by establishing correspondences between FCs based on brief and long time from the same individual. PI-FC framework maps FCs to a representation that emphasizes individual-specific patterns while retaining population-level structure as well as preserving demographic information while minimizing information loss (see Methods). Leveraging pre-trained strategies from other domains, this approach harnesses large-scale neuroimaging data to enhance individualized predictions from single scans, demonstrating robust generalization across temporal scales and diverse task conditions^46,47^. Inspired by meta-matching^48–50^, we implemented an extended variant PI-FC+, which incorporates individual phenotypic information (e.g., cognition and mental health) alongside demographic characteristics, enabling learned representations to inherently capture phenotype-relevant patterns for generalization to new datasets without fine-tuning.

**Fig 1.**
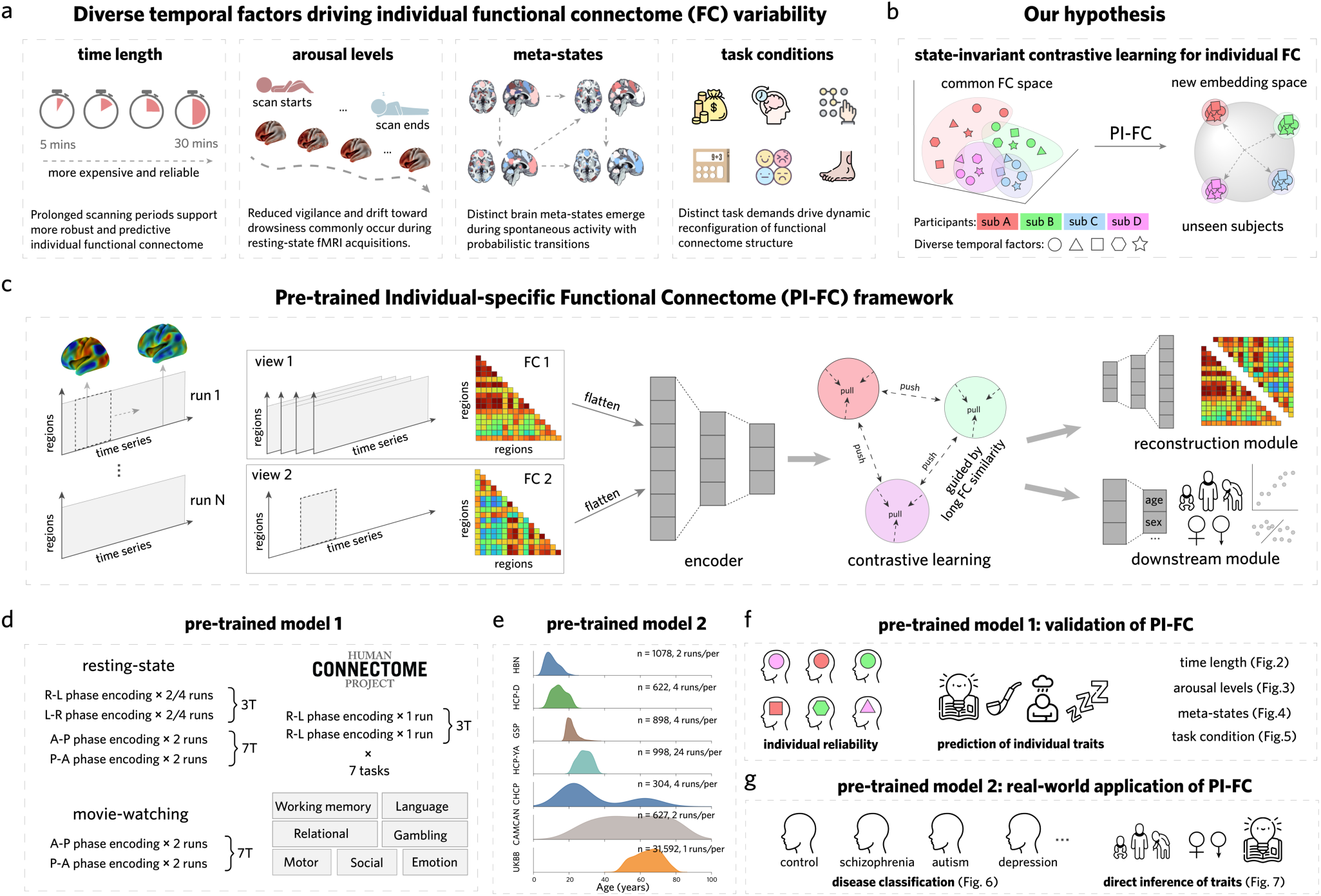
PI-FC framework for extracting stable individual-specific functional connectomes from state-dependent fluctuations. (a) Major sources of FC variability influencing individual-level reliability and clinical translation: scan duration, arousal levels, meta-states, and task conditions introduce substantial noise that masks stable individual signatures. (b) Core hypothesis: state-invariant contrastive learning transforms FC representations from a common space dominated by state-dependent fluctuations into an individual-specific embedding space that preserves stable personal characteristics. (c) PI-FC architecture employs deep contrastive learning with three complementary modules: (i) encoder for feature extraction, (ii) contrastive learning module for individual-specific optimization, and (iii) reconstruction and downstream prediction modules to preserve connectivity patterns and phenotypic information. (d) *Pre-trained model 1* is developed using densely sampled HCP-YA (Human Connectome Project Young Adult) data (489 subjects) across multiple paradigms (resting-state and seven tasks) and field strengths (3T/7T), enabling robust temporal and state-invariant learning. (e) *Pre-trained model 2* is developed leveraging large-scale multi-center datasets (> 30,000 subjects from (Healthy Brain Network; HBN), (Lifespan Human Connectome Project Development; HCP-D), (Brain Genomics Superstruct Project; GSP), HCP-YA, (Chinese Human Connectome Project; CHCP), (Cambridge Centre for Ageing and Neuroscience; CAM-CAN), (UK Biobank; UKBB)) with sequential fine- tuning and ensemble stacking strategy to enhance cross-site generalization. (f) Comprehensive validation demonstrating superior performance in individual identification, phenotypic prediction accuracy, feature stability across temporal scales (30 seconds to 15 minutesk; Fig. 2), arousal states (Fig. 3), meta-states (Fig. 4), and diverse experimental conditions (Fig. 5). (g) Real-world clinical applications including psychiatric disorder classification (schizophrenia, autism, depression, anxiety; Fig. 6) and direct inference of individual traits (age, sex, cognitive ability; Fig. 7) in independent clinical centers without site-specific retraining. Abbreviations: sub, subject.

To validate PI-FC, we implemented a two-stage approach. First, we trained *pre-trained model 1* (**Fig. 1d, left**) using data from densely sampled subjects from Human Connectome Project Young Adult (HCP-YA). The subset encompasses multiple sessions of various paradigms acquired at both 3T and 7T field strengths, with cumulative scanning durations ranging from approximately 1 hour on average to up to 4 hours per subject^4^. In an independent HCP-YA cohort, we demonstrated that the resulting PI-FC significantly improved individual identification and predictions of cognitive, mental health, and demographic phenotypes (**Fig. 1f**). Notably, these improvements were consistent across various temporal scales (from tens of seconds to hours), different brain states (including arousal states and unsupervised clustering-derived states), and diverse experimental conditions (including resting-state and seven distinct tasks spanning motor, social, language, emotion, relational, gambling, and working memory domains). Second, we developed *pre-trained model 2* (**Fig. 1e**) by leveraging multiple large-scale neuroimaging datasets totaling over 30,000 subjects^51^. Rather than aggregating data across all imaging centers, we proposed a strategy of sequentially fine-tuning models for each individual center followed by a stacking approach. This stacking strategy substantially improved generalization to previously unseen imaging centers (**Fig. 1g**), enhancing diagnostic accuracy across various psychiatric disorders while enabling direct inference of individualized phenotypic characteristics without additional training. Together, PI-FC provides a generalizable path to stable individual signatures and stronger brain–behavior links, positioning FC for more reliable clinical applications.

## Results

### PI-FC demonstrates equivalent performance using substantially shorter scan durations

PI-FC framework was designed to learn stable, inherent individual-specific FC patterns while reducing state-dependent noise across different temporal scales. We hypothesized that PI-FC could achieve high test-retest reliability and robust individualized predictions using substantially shorter scan durations compared to traditional FC approaches. To test this hypothesis, we trained *pre-trained model 1* using 489 densely sampled subjects from the HCP-YA dataset, then evaluated the model on a held-out subset of HCP-YA subjects not used in training. We simulated short-duration resting-state conditions by extracting sequential segments from complete (resting-state fMRI) rs-fMRI time series (**Fig. 2a**), generating subsets ranging from 50 frames (36 seconds) to 1,200 frames (14.4 minutes)^52,53^.

**Fig 2.**
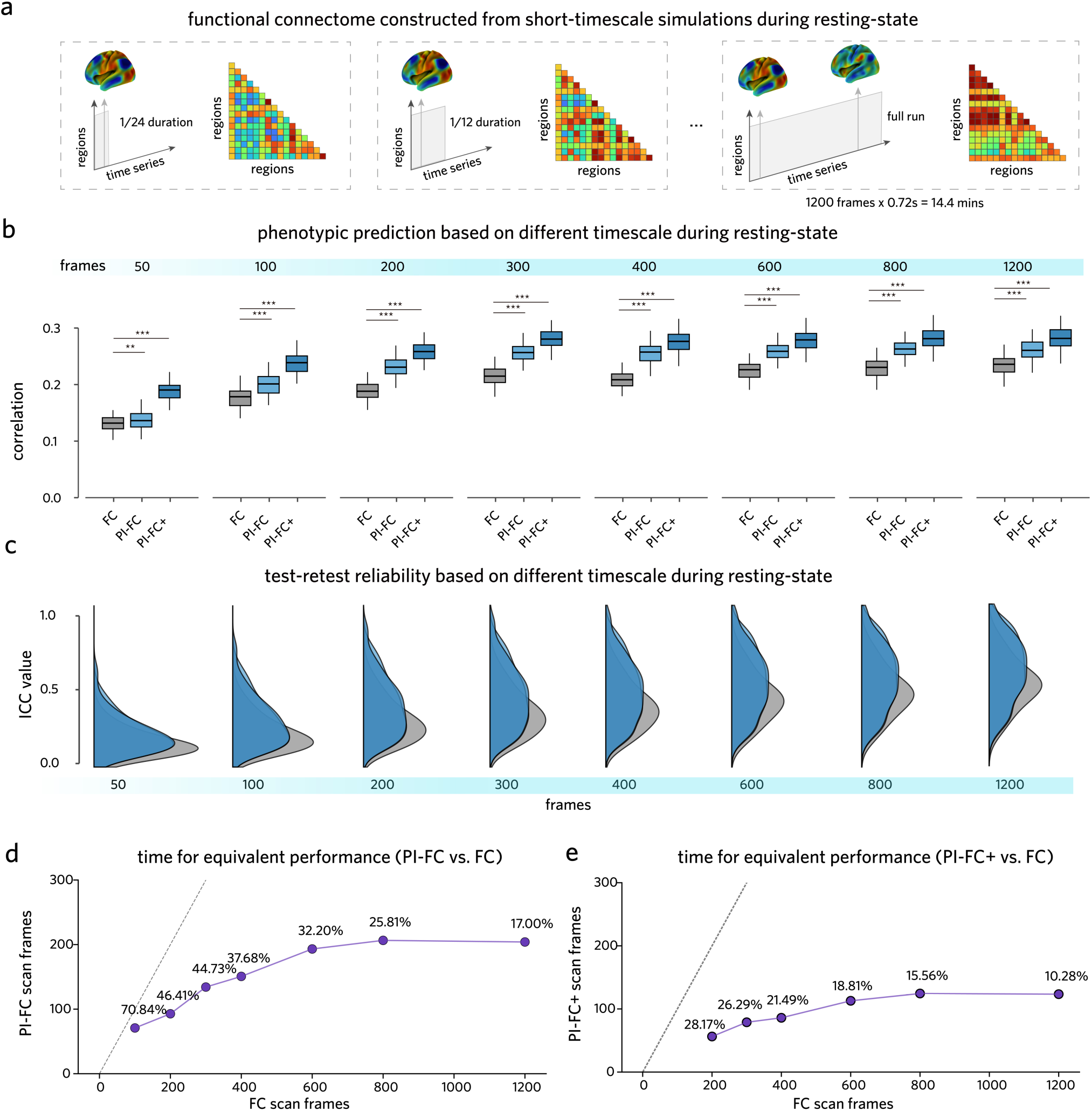
PI-FC exhibits superior predictive performance, stability, and time efficiency when applied to short-duration rs-fMRI data. (a) Schematic illustration of functional connectome construction from short- timescale simulations during resting-state, showing progressive extraction from 1/24 duration to full run (1,200 frames × 0.72s = 14.4 min). (b) Phenotypic prediction performance across varying rs-fMRI scan lengths (50 to 1,200 frames). Box plots show Pearson’s r between predicted and actual phenotypic measure (NIH toolbox total cognition composite score). Both PI-FC and PI-FC+ achieve significantly higher prediction correlations than traditional FC across nearly all scan lengths with the exception of PI-FC at 50 frames (paired t-test for most comparisons as indicated by asterisks ***, *P* < 0.001; **, *P* < 0.01). Notably, PI-FC methods maintain robust performance even with very short scan segments (e.g., 100 frames). (c) Test-retest reliability of FC features, quantified by ICC(3,1), across different scan lengths. Density plots show the distribution of ICCs across features; mean ± SD are indicated (e.g., at 1,200 frames: PI-FC 0.50 ± 0.19 vs FC 0.41 ± 0.15). Paired t-tests; BH-FDR across durations, *Q* < 0.001. (d, e) Scan time (frames) required to achieve equivalent predictive performance. For each observed accuracy value, we linearly interpolated frames as a function of performance for FC and for PI-FC (or PI-FC+), and computed—given a traditional-FC duration (x-axis)—the PI-FC (d) or PI-FC+ (e) duration (y-axis) that attains the same predictive performance. Points show the resulting equivalence mapping y(x); the dashed line denotes y = x (equal time). Labels indicate the ratio y/x (percentage of traditional FC time). Points and lines indicate point estimates from curves averaged across seeds; 95% CIs (paired bootstrap over seeds, B=10,000) are reported in the text and Supplementary Table S1 and are not plotted.

We generated functional connectomes (traditional FC, PI-FC, and PI-FC+) from these simulated short-duration data to predict 39 cognitive and mental health phenotypic measures, controlling for family structure^54^ and regressing out sex and age as confounds (see Methods). Predictive accuracy analysis across all simulated scan lengths revealed that PI-FC(+) consistently outperformed traditional FC approaches (**Fig. 2b**, cognitive performance; Supplementary Fig S1, all phenotypic measures). Using 100 independent iterations of 5-fold nested cross-validation, PI-FC and PI-FC+ achieved significantly higher phenotypic predictive performance than traditional FC at every scan duration (paired t-tests across resamples on the macro-averaged score; all *Ps* < 0.001), with the exception of PI-FC at the shortest duration (50 frames, *P* < 0.01). Notably, even with very short scan segments (e.g., 100 frames, 72 s), PI-FC and PI-FC+ maintained robust predictive performance (paired t-test, all *Ps* < 0.001). Averaged across 39 phenotypic measures, PI-FC demonstrated 19.50% improvement over traditional FC, while PI-FC+ showed even greater enhancements of 34.01% across different scan durations.

Beyond predictive accuracy, we evaluated feature stability using test-retest reliability measures across 4 runs per subject. Feature reliability was assessed using intraclass correlation coefficients (ICC, see Methods)^55,56^ calculated per edge and macro-averaged across edges from 100 independent random sub- samplings per scan length (**Fig. 2c**). PI-FC consistently yielded higher ICCs (mean ± SD: 0.13 ± 0.10 at 50 frames to 0.50 ± 0.19 at 1200 frames) compared to traditional FC (0.08 ± 0.08 to 0.41 ± 0.15; paired t-test; BH-FDR across durations, *Q* < 0.001), demonstrating enhanced measurement reliability across all temporal scales.

To quantify PI-FC’s scan time efficiency, we determined the scan time required for PI-FC to achieve predictive performance equivalent to that of traditional FC (**Fig. 2d-e**). The analysis compared traditional FC scan time (x-axis) against the time required by PI-FC (**Fig. 2d**) or PI-FC+ (**Fig. 2e**) to reach the same macro-averaged predictive performance. At performance equivalence to FC at 1,200 frames (14.4 min), PI-FC required 204 frames (2.45 min; 17.0% of FC time, 95% CI 16.5–17.9%), whereas PI-FC+ required 123 frames (1.48 min; 10.3%, 95% CI 9.5–11.2%). These results indicate that PI-FC effectively captures individual-specific information from brief scanning durations.

### PI-FC maintains robust performance across different arousal states

While we observed substantial improvements in phenotypic prediction with PI-FC at shorter durations, gains plateaued between 800 and 1,200 frames, suggesting that factors beyond scan length might influence FC reliability. Prior work indicates that vigilance systematically declines during extended resting-state acquisitions, potentially confounding individual-specific connectivity^20,57,58^. To examine whether PI-FC’s advantages persist across different arousal states, we analyzed resting-state fMRI data from independent test subjects, using the initial 400 frames as a proxy for high-vigilance states and the terminal 400 frames as a proxy for low-vigilance states (**Fig. 3a**).

**Fig 3.**
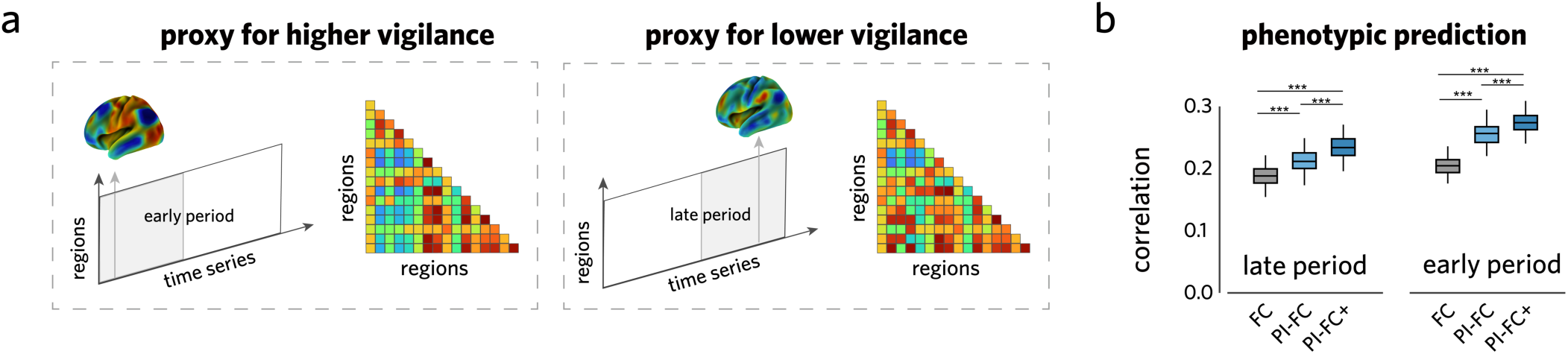
PI-FC maintains superior performance across different arousal states. (a) Schematic of functional connectome construction from resting-state data. The initial 400 frames were used as a proxy for high- vigilance states, and the terminal 400 frames as a proxy for low-vigilance states. Detailed procedures are described in the Methods. (b) Phenotypic prediction across arousal states. Box plots show Pearson’r between predicted and actual phenotypic measure (NIH toolbox total cognition composite score). Both PI-FC and PI- FC+ achieve significantly higher prediction correlations than traditional FC across both early and late periods (paired t-test, ***,*P* < 0.001). Notably, PI-FC applied to late-period data achieves performance comparable to traditional FC from early periods, demonstrating effective compensation for vigilance-related performance degradation.

Phenotypic prediction analysis showed that PI-FC and PI-FC+ outperformed traditional FC across both high- and low-vigilance states (**Fig. 3b**, cognitive performance; Supplementary Fig. S2, all phenotypic measures). Notably, whereas traditional FC degraded during likely low vigilance states periods, both PI- FC and PI-FC+ variants demonstrated consistent superiority across both early and late scanning periods (paired t-test, all *Ps* < 0.001). Critically, PI-FC applied to low vigilance data achieved prediction performance comparable to traditional FC during high-vigilance states, indicating effective compensation for vigilance-related performance degradation. These findings demonstrate that PI-FC mitigates confounding effects of arousal fluctuations, a major source of variance in clinical fMRI acquisitions.

### PI-FC captures stable individual signatures across distinct brain states

Building on PI-FC’s robustness across scan durations and vigilance states, we next examined whether these advantages extend to spontaneous brain state transitions during resting-state scanning. Previous work has shown that rs-fMRI exhibits dynamic fluctuations between distinct connectivity patterns, representing different meta-states of brain activity^59,60^. To test PI-FC’s stability across these naturally occurring state variations, we identified predominant meta-states and assessed whether PI-FC could transform state-specific connectivity patterns into consistent individual signatures (**Fig. 4a**). Using a sliding window approach with 50 TR windows (36-second segments, no overlap) and k-means clustering (see Methods), we identified 4 meta-states from the trained HCP-YA cohort with distinct occurrence frequencies (state 1: 40.0%, state 2: 28.6%, state 3: 16.4%, and state 4: 15.0%). For comparison, each individual’s “long FC” represented their stable trait-like connectivity profile averaged across all scanning sessions.

**Fig 4.**
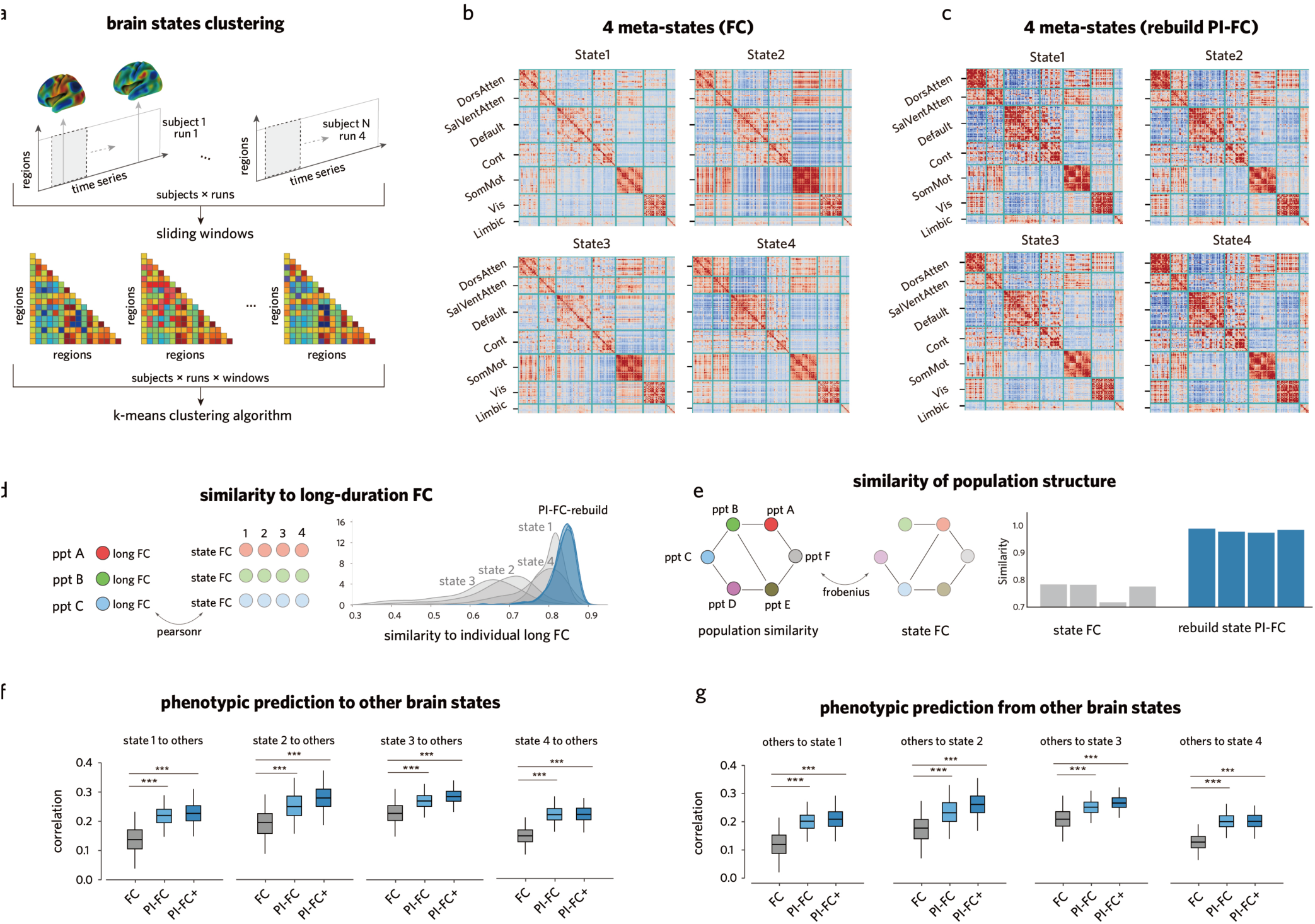
PI-FC mitigates state-dependent fluctuations to capture stable, individual-specific functional connectomes. (a) Schematic illustration of brain state clustering methodology using sliding windows and k- means clustering to identify 4 meta-states from rs-fMRI. (b) Traditional FC matrices for 4 meta-states (State 1-4, occurrence frequencies: 15.0%-40.0%), show distinct patterns across meta-states. (c) rebuild PI-FCs, exhibit reduced state-specific variations and enhanced similarity across all meta-states. (d) rebuild PI-FCs show higher similarity to individual long FC. Density plots demonstrate higher Pearson’s r for PI-FC (blue distribution) compared to original state FCs (grey distribution; paired t-test *P* < 0.001). (e) PI-FC enhances correspondence with phenotypic structure. Similarity scores (Frobenius norm) show rebuild PI-FCs correspond significantly better to behavioral phenotypic similarity matrices than original state FCs (paired t- test; *P* < 0.001). (f, g) Cross-state phenotype prediction performance (NIH toolbox total cognition composite score). Box plots show Pearson’s *r*. PI-FC (mean ± SD = 0.24 ± 0.04) and PI-FC+ (mean ± SD = 0.25 ± 0.04) exceed traditional FC (mean ± SD = 0.17 ± 0.06; paired t-test, *P* < 0.001) when (f) generalizing from each state to all other states, and (g) generalizing from all other states to each specific state.

Visual inspection revealed marked differences between traditional and PI-FC approaches. Although time series were subjected to global signal regression (GSR)^61–63^, traditional state-specific FCs still showed distinguishable patterns that varied substantially across meta-states (**Fig. 4b**). For more intuitive demonstration, we applied a nonlinear decoder to map embeddings back into FC space, yielding rebuild PI-FC (see Methods) that enabled direct comparison within the FC space and demonstrated attenuated meta-state effects (**Fig. 4c**). This observation was confirmed quantitatively: rebuild PI-FCs showed consistently higher similarity to individual long FCs across all states (state 1: mean *r* = 0.84, state 2: mean *r* = 0.84, state 3: mean *r* = 0.83, state 4: mean *r* = 0.83; *Ps* < 0.001) compared to original state FCs (state 1: mean *r* = 0.80, state 2: mean *r* = 0.68, state 3: mean *r* = 0.62, state 4: mean *r* = 0.77; *Ps* < 0.001) (**Fig. 4d**). Additionally,rebuild PI-FCs improved correspondence between FC patterns and phenotypic similarity structures across all states (**Fig. 4e**).

Cross-state generalization further highlighted PI-FC’s advantage. We performed bidirectional validation where models were trained on individual meta-states and tested across all remaining states for cognitive performance prediction (see Methods). Both PI-FC and PI-FC+ outperformed traditional FC in predicting cognition (paired t-test, all *Ps* < 0.001; Supplementary Fig. S3, all phenotypic measures), corresponding to relative improvements of 38% and 56%, respectively (**Fig. 4f**). In the reverse validation, where models were trained on multiple states and tested on individual held-out states (**Fig. 4g**), PI-FC methods maintained their performance advantage, confirming robustness to spontaneous meta-state variation.

### PI-FC captures task-invariant individual signatures enabling robust cross-paradigm generalization

Having demonstrated PI-FC’s robustness to temporal and state-dependent fluctuations, we next tested whether PI-FC preserves individual-specific signatures across diverse cognitive tasks. This is a critical test of PI-FC’s utility, as successful cross-paradigm integration would enable combining datasets from different experimental designs, increasing effective sample sizes and statistical power for phenotypic analyses.

We randomly selected 200 subjects from the independent HCP-YA subjects and repeated this process 100 times for analysis: for each subject, we extracted 100-frame (72s) segments from resting-state and seven distinct task paradigms—emotion, language, motor, relational, social cognition, working-memory, and gambling—spanning diverse cognitive demands with multiple runs per paradigm. Traditional FC achieved reasonable within-task identification accuracy (62.8 ± 16.0%) but showed severely degraded performance cross-task (7.3 ± 8.8%; **Fig. 5a**). In contrast, PI-FC both improved within-task accuracy (80.7 ± 5.5%) and critically maintained substantial cross-task identification accuracy (48.2 ± 16.9%; paired t-test, all *Ps* < 0.001; **Fig. 5b**), revealing a robust task-invariant brain fingerprint.

**Fig 5.**
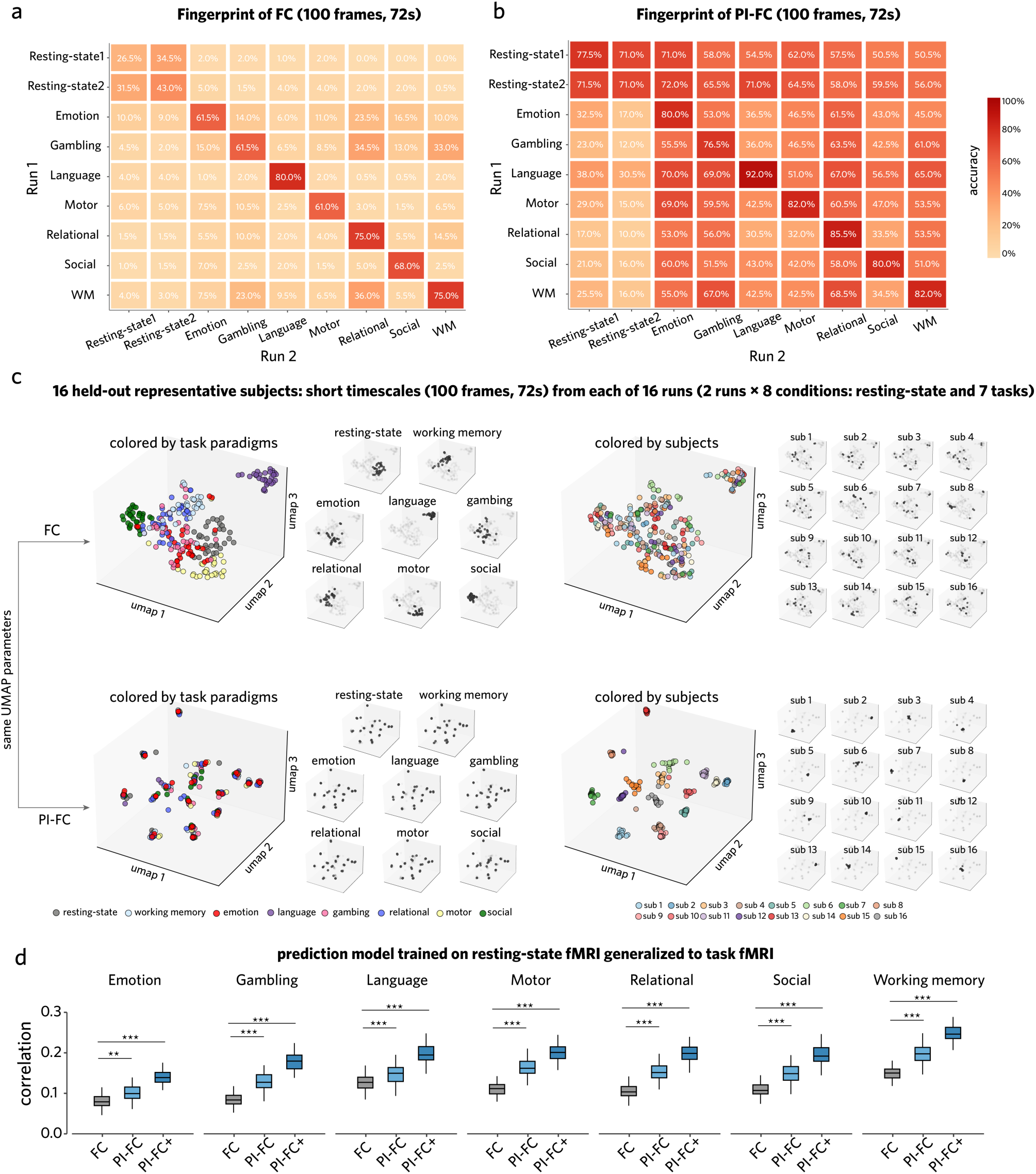
PI-FC enhances individual-specific representations for superior cross-task generalization. (a-b) Individual fingerprinting accuracy matrices using 100-frame (72s) segments across resting-state and seven task paradigms. (a) Traditional FC shows moderate within-task identification (diagonal: 62.8% ± 16.0%) but poor cross-task accuracy (off-diagonal: 7.3% ± 8.8%). (b) PI-FC improves both within-task (80.7% ± 5.5%) and cross-task identification (48.2% ± 16.9%; *P* < 0.001). (c) UMAP embeddings of FC samples from 16 representative subjects across eight paradigms. When labeled by paradigms (left panels), traditional FC forms task-locked clusters while PI-FC shows intermixing across tasks. When labeled by subjects (right panels), traditional FC scatters broadly whereas PI-FC forms tight subject-centered clusters. PI-FC achieves higher individual-identity clustering coefficients (0.93 ± 0.28) and lower task-clustering coefficients (-0.69 ± 0.24) compared to traditional FC (*P* < 0.001). (d) Cross-task cognitive prediction performance (NIH toolbox total cognition composite score). Models trained on resting-state data and tested on task data show PI-FC and PI- FC+ consistently outperform traditional FC across all tasks (all *Ps* < 0.001).

UMAP^64^ visualization of 16 representative subjects illustrated how PI-FC disentangles task-specific from individual-specific information (**Fig. 5c**). When labeled by paradigms, traditional FC formed distinct task- locked clusters, whereas PI-FC samples intermixed across tasks. When labeled by individual, the pattern reversed: traditional FC scattered broadly, while PI-FC formed tight, subject-centered clusters. Quantitatively, PI-FC achieved higher individual-identity clustering coefficients (0.93 ± 0.28) and lower task-clustering coefficients (-0.69 ± 0.24) than traditional FC (*P* < 0.001), confirming preferential capture of stable individual characteristics over different task demands.

This enhanced individual-specificity translated into superior cross-paradigm prediction. Models trained exclusively on resting-state data and tested on task data without retraining—simulating realistic cross- paradigm transfer—showed that PI-FC and PI-FC+ consistently outperformed traditional FC across all task conditions for cognitive performance (paired t-test, all *Ps* < 0.001; **Fig. 5d**). Similar patterns were observed when averaged across all 39 phenotypic measures tested (Supplementary Fig. S4). On mixed- paradigm datasets combining resting-state and task data with paradigms randomly selected for the training and test sets, PI-FC and PI-FC+ significantly exceeded traditional FC performance (paired t-test, all *Ps* < 0.001).

### Stacked multi-source pre-training enhances diagnostic accuracy across independent centers for psychiatric classification

Functional connectomes show promise as psychiatric biomarkers but face key barriers to clinical translation, including patient intolerance to prolonged scanning and substantial cross-center heterogeneity in imaging protocols^65–67^. To address these challenges, we developed *pre-trained model 2* by sequentially fine-tuning our HCP-YA foundation model (*pre-trained model 1*) across multiple large-scale datasets (HCP-D^68^, Cam-CAN^9^, CHCP^69^, UKBB^51^, GSP^70^, HBN^71^; see Methods), generating center-specific models while preserving individual-specific representations. We then implemented a stacked ensemble approach that averages prediction probabilities from all non-overlapping source models for each target cohort, leveraging complementary strengths while minimizing dataset-specific biases.

We evaluated this PI-FC+ (stacking) approach against traditional FC and the single pre-trained PI-FC+ (HCP-YA) model across 4 independent psychiatric cohorts: schizophrenia (HCP-EP)^72^, autism spectrum disorder (ABIDE)^11^, generalized anxiety disorder (GAD), depression (SRPBS)^73^. Statistical significance was assessed through Wilcoxon signed-rank tests with Holm-Bonferroni correction. PI-FC+ (stacking) consistently outperformed both traditional FC and single pre-trained models across these psychiatric conditions (**Fig. 6a-d**: schizophrenia, autism, GAD and depression). The largest gains were observed for GAD classification, achieving 74.9% ± 3.7% balanced accuracy (bACC) and 84.2% ± 3.2% under the ROC curve (AUC)—representing 18.6% and 31.0% improvements over traditional FC, respectively (*P* = 0.002/0.002 for bACC/AUC). Significant improvements were also observed in schizophrenia (HCP-EP: *P* = 0.002/0.005 for bACC/AUC), autism spectrum disorder (ABIDE: *P* = 0.002/0.002 for bACC/AUC), and depression (SRPBS: *P* = 0.0039 for bACC), with comprehensive metrics detailed in **Fig. 6** and Supplementary Tables S2-S5.

**Fig 6.**
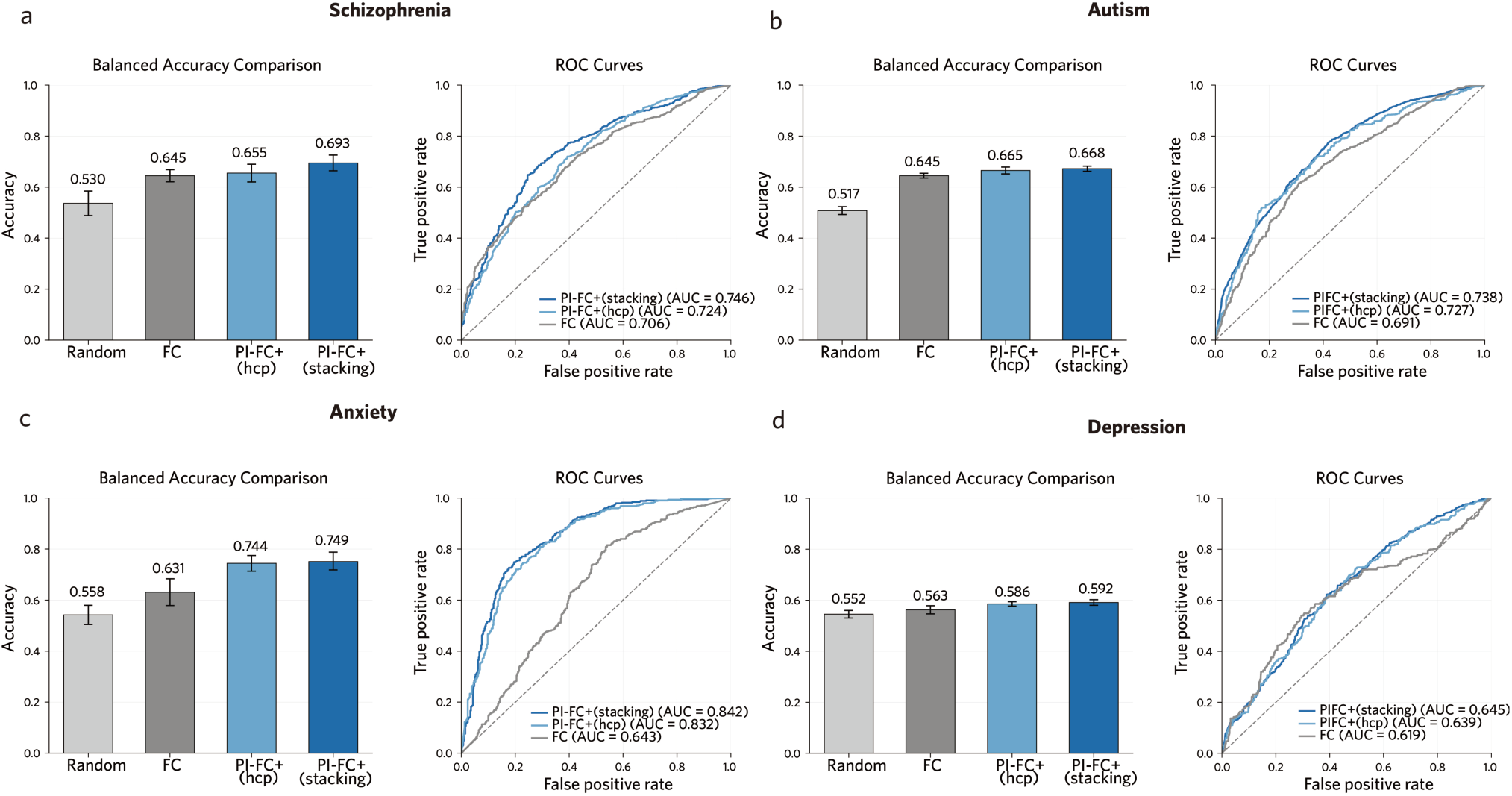
Multi-source pre-training and stacking strategy enhance classification robustness and generalization of PI-FC+ across multi-disease clinical cohorts. Classification performance comparison between traditional FC, PI-FC+ (HCP-YA) single pre-trained model, and PI-FC+ (stacking) multi-source ensemble for distinguishing patients from healthy controls across 4 independent psychiatric cohorts: (a) HCP-EP schizophrenia [Traditional FC: 64.5% ± 2.5% bACC, 70.6% ± 2.2% AUC; PI-FC+ (HCP-YA): 65.5% ± 3.7% bACC, 72.4% ± 2.4% AUC; PI-FC+ (stacking): 69.3% ± 3.2% bACC, 74.6% ± 2.4% AUC], (b) ABIDE autism spectrum disorder [Traditional FC: 64.5% ± 1.0% bACC, 69.1% ± 0.7% AUC; PI-FC+ (HCP-YA): 66.5% ± 1.4% bACC, 72.7% ± 2.0% AUC; PI-FC+ (stacking): 66.8% ± 1.1% bACC, 73.8% ± 0.6% AUC], (c) GAD generalized anxiety disorder [Traditional FC: 63.1% ± 5.5% bACC, 64.3% ± 4.7% AUC; PI-FC+ (HCP-YA): 74.4% ± 3.2% bACC, 83.2% ± 2.7% AUC; PI-FC+ (stacking): 74.9% ± 3.7% bACC, 84.2% ± 3.2% AUC], and (d) SRPBS depression [Traditional FC: 56.3% ± 1.7% bACC, 61.9% ± 4.1% AUC; PI-FC+ (HCP-YA): 58.6% ± 0.9% bACC, 63.9% ± 0.9% AUC; PI-FC+ (stacking): 59.2% ± 1.5% bACC, 64.5% ± 1.4% AUC]. Left panels display bACC with error bars representing standard deviation; right panels show ROC curves with corresponding AUC values. Statistical significance was assessed using Wilcoxon signed-rank tests with Holm-Bonferroni correction. Detailed statistical comparisons and additional performance metrics are provided in Supplementary Tables S6-S7.

These findings demonstrate that stacking multi-source, center-specific models enhances the robustness and generalizability of individual-specific connectivity features, improving psychiatric classification across independent clinical centers.

### PI-FC+ (stacking) enables robust inference of individual biological and cognitive phenotypes

Having demonstrated superior diagnostic performance, we next evaluated whether PI-FC+(stacking) could directly infer individual phenotypes in completely independent clinical centers without additional training. This capability represents a significant advance for clinical translation, enabling immediate deployment of pre-trained models for personalized brain function assessment. We tested age, biological sex, and cognitive ability across 4 independent clinical cohorts (ABIDE, HCP-EP, SRPBS, GAD) that were entirely separate from pre-training datasets and non-overlaping with source models used in stacking.

PI-FC+ (stacking) demonstrated robust direct inference capabilities across all tested phenotypes (**Fig. 7a-b**). For age prediction (**Fig. 7a**), the model achieved significant correlations between predicted and actual age across all 4 datasets (ABIDE: *r* = 0.588; HCP-EP: *r* = 0.265; SRPBS: *r* = 0.553; GAD: *r* = 0.488; all *Ps* < 0.01), although the relatively narrow age span in HCP-EP likely attenuated the correlation due to range restriction. Performance remained consistent within both healthy control and patient subgroups, demonstrating robustness across clinical heterogeneity (**Fig. 7a)**. For sex classification, PI-FC+ achieved high discrimination accuracy across cohorts, with AUC ranging from 0.786 (SRPBS) to 0.953 (GAD) (**Fig. 7b**). For cognitive ability, PI-FC+ (stacking) achieved significant predictions in the two cohorts with available data: ABIDE (*r* = 0.194, *P* < 0.001) and HCP-EP (*r* = 0.406, *P* < 0.001).

**Fig 7.**
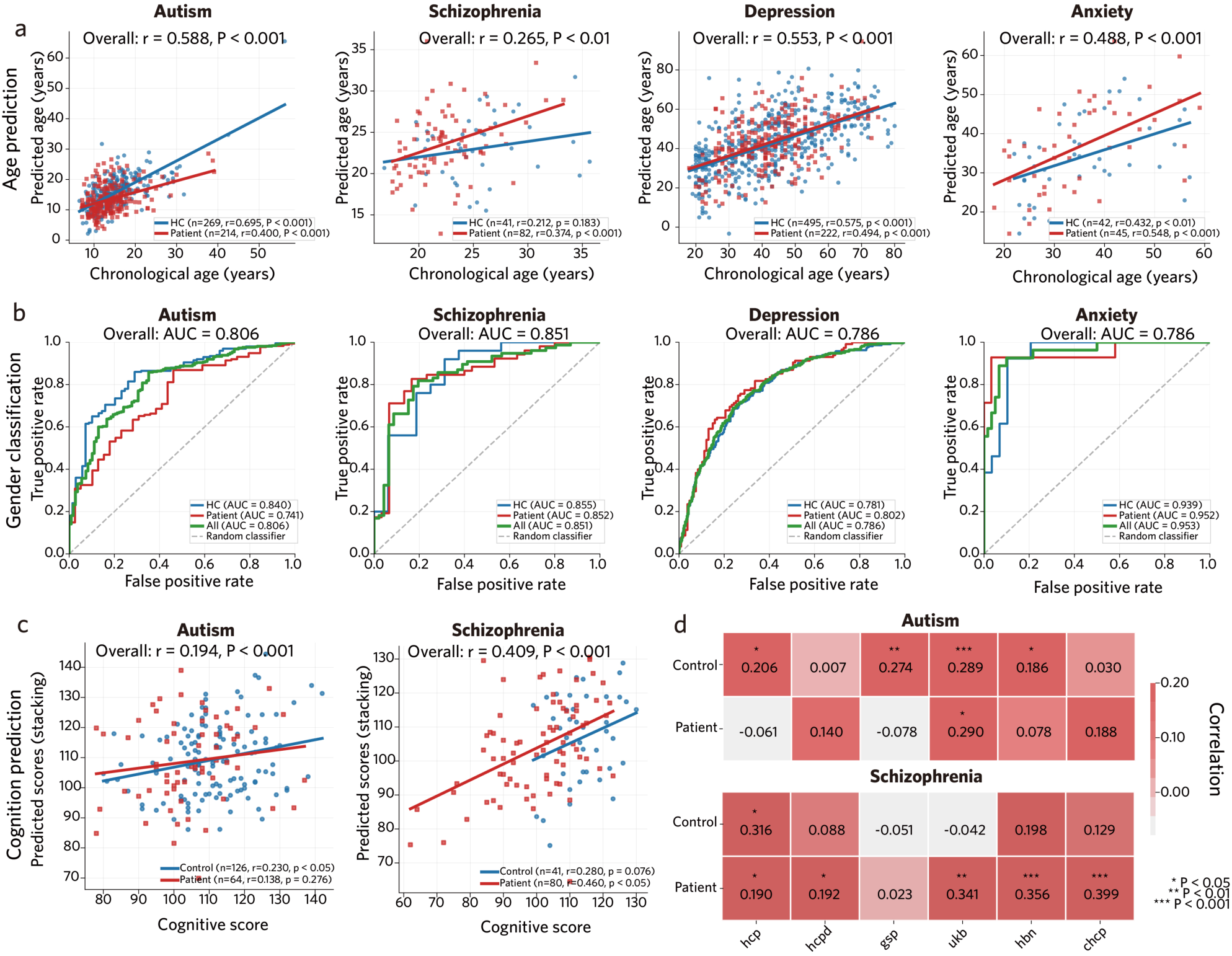
PI-FC+ (stacking) enables direct inference of individual biological and cognitive phenotypes across independent clinical centers. Performance of the PI-FC+ (stacking) ensemble model in directly inferring individual age, biological sex, and cognitive ability across multiple independent clinical and healthy mixed cohorts. Further analyses dissect the cognitive ability prediction performance of individual pre-trained models contributing to the stacking ensemble. (a) Age prediction using PI-FC+ (stacking). Scatter plots show significant correlations (Pearson’s r) between predicted age and actual age in the ABIDE (r = 0.588, *P* < 0.001), HCP-EP (r = 0.265, *P* < 0.01), SRPBS (r = 0.553, *P* < 0.001), and GAD (r = 0.488, *P* < 0.001) cohorts. Correlations within healthy control and patient subgroups are also displayed with different colors. (b) Sex classification performance of PI-FC+ (stacking). ROC curves show high AUC values for distinguishing biological sex in the ABIDE (AUC = 0.806), HCP-EP (AUC = 0.851), SRPBS (AUC = 0.786), and GAD (AUC = 0.953) datasets. (c) Cognitive ability prediction using PI-FC+ (stacking). Scatter plots show significant correlations (Pearson’s r) between predicted and observed cognitive scores in the ABIDE (*r* = 0.194, *P* < 0.001) and HCP-EP (*r* = 0.406, *P* < 0.001) datasets. (d) Cognitive ability prediction performance (Pearson’s r) of individual PI-FC+ source models (pre-trained on HCP-YA, HCP-D, GSP, UKBB, HBN, CHCP) on the ABIDE (upper panel) and HCP-EP (lower panel) test sets. Performance is shown for the total sample, healthy controls, and respective patient subgroups (Autism for ABIDE; Schizophrenia for HCP-EP). Asterisks denote statistical significance (* *P* < 0.05, ** *P* < 0.01, *** *P* < 0.001; one-sided test). For example, In the ABIDE autism subgroup, models trained on UKBB (*r* = 0.29, *P* < 0.05) showed stronger predictive correlations. In the HCP-EP schizophrenia subgroup, models trained on CHCP (*r* = 0.399, *P* < 0.001), HBN (*r* = 0.356, *P* < 0.001), and UKBB (*r* = 0.341, *P* < 0.001) demonstrated notable predictive ability.

To understand the ensemble advantage, we examined individual model contributions to cognitive prediction (**Fig. 7d**). Different pre-training datasets showed varying effectiveness across populations: for example, in ABIDE autism group, models trained on UKBB (*r* = 0.29, *P* < 0.05) contributed most; while in HCP-EP schizophrenia group, CHCP (*r* = 0.399, *P* < 0.001), HBN (*r* = 0.356, *P* < 0.001), and UK Biobank (*r* = 0.341, *P* < 0.001) models were most effective. This population-specific variation highlights the value of ensemble stacking, which leverages complementary strengths of multiple models trained on diverse datasets to achieve more robust and accurate phenotype inference than any single model alone.

These findings establish PI-FC+ (stacking) as a generalizable framework for direct individual phenotype inference across unseen clinical centers, enabling personalized brain function assessment without site- specific model retraining.

## Discussion

Functional connectomes contain high-dimensional data that encode rich information about both individual traits and state-dependent fluctuations, with the latter containing uncontrollable variability that reduces reliability and limits clinical translation^18–23,36,37^. In this work, we present PI-FC, a pre-trained deep learning framework that addresses fundamental state-dependent confounding factors in functional connectomes. PI-FC employs state-invariant contrastive learning to extract stable, trait-related signatures while suppressing state-dependent confounders, supporting reliable estimation of individual-level functional characteristics that generalize to unseen subjects. Validation using p*re-trained model 1* on independent HCP-YA cohorts showed that PI-FC achieved comparable phenotypic prediction with 17% of FC scan time (1,200 frames; equivalence analysis), maintained robust performance across diverse meta-states, and generalized across different experimental paradigms. The multi-source ensemble strategy (PI- FC+ stacking), leveraging over 30,000 subjects across 8 independent datasets for *pre-trained model 2*, further enhanced diagnostic accuracy across multiple psychiatric disorders while enabling direct inference of individual traits without additional training on new clinical datasets. These findings provide evidence that PI-FC overcomes fundamental barriers to neuroimaging reliability and establishes functional connectomics as a viable foundation for clinical translation.

Two lines of prior neuroscience evidence guided its conceptual development: First, seminal fingerprinting studies showed that resting-state functional connectome can accurately identify individuals across scanning sessions, days, and even years, with recent deep learning approaches showing that as little as 20 seconds of data contains sufficient individual-specific information for reliable identification^14,39,40^. Second, previous work revealed that resting-state and task-based paradigms contain complementary information while sharing core individual differences, suggesting the existence of a stable, state-invariant FC architecture that optimally predicts phenotypic measures^19,36–38,74,75^. However, these supervised approaches require extensive within-subject data across multiple sessions and do not generalize zero-shot to entirely new individuals—a critical limitation for clinical deployment. PI-FC operationalizes state invariance by coupling a contrastive encoder with (i) a structure-preserving constraint that maintains population-level similarity geometry between embedding and FC spaces, and (ii) a decoder that explicitly recovers long FC patterns from brief FC inputs. This combination regularizes representations toward trait-like identity while preventing dispersion of subject embeddings that can attenuate phenotype-relevant differences, enabling direct application to unseen individuals from a single scanning run.

Empirical evaluation confirmed that PI-FC effectively mitigates the influence of vigilance- and task-related fluctuations on FC-based prediction of behavior. Longer scan durations generally improve phenotypic prediction^18,22,26^, and early scan periods (reflecting higher vigilance states) outperform later periods when subjects may experience fatigue or reduced alertness. This temporal degradation is problematic given that approximately 30% of subjects experience difficulty maintaining stable wakefulness beyond 3 minutes during resting-state scanning, leading to progressive changes in connectivity patterns that introduce state-dependent variability^20–22^. Similarly, cross-task generalization with traditional FC is poor, reflecting the substantial state-dependent variance that confounds individual-specific information. PI-FC mitigates these state-dependent influences by enabling robust cross-paradigm generalization. This capability addresses a critical need in the field for methods that can harmonize data across different scanning protocols, paradigms, and temporal scales. While global signal regression is commonly employed to reduce arousal-related and other state-dependent confounds^61,63^, we observed further reduction of state-related differences after PI-FC reconstruction relative to GSR alone, consistent with extraction of trait-like connectivity patterns.

From a clinical translation perspective, our ensemble strategy provides a practical approach to address scanner and site-heterogeneity challenges in developing neuroimaging biomarkers. Considering the practical costs of clinical neuroimaging, our framework provides an approach to construct stable individual-specific functional connectivity representations using reduced resting-state scan durations and by integrating information across different task paradigms, which may have important implications for real-world applications. Importantly, we empirically evaluated sequential fine-tuning followed by stacking as well as training a single unified model on aggregated multi-center data—the approach typically adopted by current neuroimaging foundation models^46,47^. However, the straightforward data amalgamation approach may be limited by imbalance and heterogeneity across datasets, underperforming compared to single-center PI-FC+ (HCP-YA); in contrast, our stacking strategy^76,77^ consistently demonstrated advantages over single-center PI-FC+ (HCP-YA) in psychiatric disease classification tasks across independent centers. The results could provide insights for other foundation model development efforts, suggesting that sequential fine-tuning plus ensemble integration may offer advantages over direct data amalgamation. Furthermore, our framework incorporates meta-matching principles^48,49^ that facilitate direct inference of new traits in independent datasets, exemplified by zero-shot prediction of age, biological sex, and cognitive abilities in completely new clinical populations without requiring additional training data.

Several limitations warrant consideration. Due to computational complexity from dense temporal sampling across large-scale datasets, we employed a relatively coarse ∼200 parcellation that may not capture fine-grained connectivity patterns potentially enhancing individual specificity. Future work should investigate higher spatial resolution approaches. Our current framework for direct inference focuses on core phenotypes (age, biological sex, basic cognitive measures); expansion to diverse psychiatric symptoms, personality traits, and complex behavioral phenotypes would broaden clinical utility. If pretraining datasets encompass diverse clinical populations, the framework may enable direct disease inference without disease-specific training. Additionally, we employed Pearson correlation-based functional connectome, representing one approach among many possible connectivity measures. Recent work has benchmarked over 200 pairwise correlation methods, suggesting that integrating diverse connectivity measures into a unified model could potentially enhance representational capacity^6^. Finally, fairness and generalizability warrant study as cohorts can be imbalanced across age, sex, and site; and computational cost/inference latency should be reduced to facilitate deployment in routine workflows^78–80^. The PI-FC framework is compatible with different modalities and extends beyond fMRI to other neuroimaging modalities including EEG^81^, MEG^82,83^, and multi-modal integration analyses^84,85^.

In conclusion, PI-FC learns individual-specific traits while suppressing state-dependent information in functional connectome analyses. By achieving comparable performance with shorter scan durations and improving reliability across diverse brain states, this approach shows potential for addressing barriers in clinical translation of functional connectome.

## Methods

### Datasets and Subjects Overview

We implemented a hierarchical multi-dataset strategy encompassing 36,119 subjects from 8 independent datasets (Table 1). This comprehensive design employed two complementary pre-training approaches: *pre-trained model 1* utilized high-quality HCP-YA dataset to establish foundational brain-behavior relationships under controlled acquisition conditions, while *pre-trained model 2* incorporated the full multi-dataset collection (n = 36,119 across 8 datasets) to enhance generalizability across diverse populations, age ranges (5–87 years), and acquisition protocols. Clinical validation was performed on 4 independent patient cohorts (n = 1431) spanning neurodevelopmental and psychiatric conditions.

**Table 1.** Overview of datasets used for PI-FC framework development and validation.

| Dataset | N | Age (years) | Population | Scanner | Purpose |
| --- | --- | --- | --- | --- | --- |
| HCP-YA<sup>4</sup> | 998 | 22-35 | Healthy (US) | 3T/7T Siemens | Pre-training |
| HCP-D<sup>68</sup> | 622 | 5-21 | Healthy (US) | 3T Siemens Prisma | Pre-training |
| UKBB<sup>51</sup> | 31,592 | 44-82 | General (UK) | 3T Siemens Skyra | Pre-training |
| Cam-CAN<sup>9</sup> | 627 | 18-87 | Healthy (UK) | 3T Siemens Trio | Pre-training |
| GSP<sup>70</sup> | 898 | 18-35 | Healthy (US) | 3T Siemens Trio | Pre-training |
| HBN<sup>71</sup> | 1,078 | 5-21 | Pediatric (US) | 3T Siemens Trio | Pre-training |
| CHCP<sup>69</sup> | 304 | 18-79 | Healthy (China) | 3T Siemens Prisma | Pre-training |
| HCP-EP<sup>72</sup> | 134 | 16-35 | Psychosis (US) | 3T Siemens Prisma | Pre-training/Clinical |
| ABIDE<sup>11</sup> | 483 | 7-64 | Autism<br>(Multi-site; US) | 3T Multi-vendor | Clinical |
| SRPBS<sup>73</sup> | 727 | 18-80 | Depression<br>(Multi-site; Japan) | 3T Multi-vendor | Clinical |
| GAD | 87 | 18-59 | Anxiety (China) | 3T Siemens Prisma | Clinical |

### Pre-training Datasets

#### Human Connectome Project Young Adult (HCP-YA)

The HCP S1200 release provided neuroimaging and behavioral data from 1,094 healthy young adults (age range: 5–21 years, mean age: 28.7 ± 3.7 years; females: 531 (53.2%)), all screened to exclude neurological and psychiatric disorders. The primary 3T dataset was acquired using gradient-echo EPI sequences (TR = 720 ms, TE = 33.1 ms, flip angle = 52°, FOV = 208 × 180 mm, 72 axial slices). Resting-state fMRI consisted of two scanning sessions (resting-state 1 and resting-state 2) on separate days, each with LR and RL phase-encoding directions, comprising 1,200 frames over 14.4 minutes. The final sample included 998 subjects who completed all 4 resting-state runs. Task-based fMRI included working memory, motor, gambling, emotion, language, relational, and social tasks across two scanning days. A subset of 187 subjects underwent complementary 7T protocols (TR = 1000 ms, TE = 22.2 ms, flip angle = 45°, FOV = 208 × 208 mm, 85 axial slices). Movie-watching fMRI protocols included 4 runs with subjects viewing approximately 60 minutes of movie clips with 20- second rest periods. Resting and movie data underwent sICA-FIX preprocessing to minimize artifacts, and task data was pre-processed by HCP minimal preprocessing pipeline.

### HCP-Development (HCP-D)

The HCP-D 2.0 Release adapted HCP protocols for developmental populations, providing preprocessed rs-fMRI data from 622 typically developing subjects (age range: 5– 21 years, mean age: 14.7 ± 3.9 years; females: 336 (54.0%)) in the United States. Data were acquired using 3T Siemens Prisma scanner with gradient-echo EPI sequences (TR = 800 ms, TE = 37 ms, 2 mm isotropic voxels). The rs-fMRI protocol consisted of two scanning sessions (resting-state1 and resting- state2), each comprising two runs of 478 frames (6.4 minutes each), alternating between PA and AP phase encoding directions. Preprocessed data from the sICA-FIX pipeline were utilized.

### UK Biobank (UKBB)

We incorporated rs-fMRI data from 31,592 UK Biobank subjects (age range: 44 — 82 years, mean age: 64.1 ± 7.7 years; females: 16,448 (52.1%)) who underwent imaging between 2014 and 2022. Data were acquired at 4 dedicated imaging centers using harmonized 3T Siemens Skyra scanners. Each subject contributed a single 6-minute rs-fMRI run (490 volumes) with gradient-echo EPI sequence (TR = 735 ms, TE = 39 ms, flip angle = 52°, 2.4 mm isotropic voxels, FOV = 88×88×64). The centrally preprocessed data included motion correction, grand-mean intensity normalization, distortion correction, and sICA-FIX artifact removal, registered to MNI152 2mm standard space. Age, sex, and cognitive/emotional measures from Instance 0 and Instance 2 visits were utilized for PI-FC+ model training.

### Cambridge Center for Aging and Neuroscience (Cam-CAN)

We utilized rs-fMRI data from 627 healthy adult subjects (ages 18–87, mean age: 54.1 ± 18.5 years; females: 315 (50.2%)) from this large-scale cross-sectional lifespan study designed to characterize age-related changes in cognition and brain function. Data was acquired at MRC Cognition and Brain Sciences Unit, Cambridge, UK, using a 3T Siemens TIM Trio scanner with 32-channel head coil. Two resting-state fMRI runs per subject were acquired with subjects resting with eyes closed (TR = 1970 ms, TE = 30 ms, flip angle = 78°, 32 descending slices, slice thickness = 3.7 mm with 20% gap, voxel size = 3×3×4.44 mm, FOV = 192×192 mm, 261 volumes per run, about 8 minutes 40 seconds duration). Age and sex were utilized in Cam-CAN specific PI-FC+ model training.

### Brain Genomics Superstruct Project (GSP)

Data from 898 subjects were selected from this repository of neuroimaging, behavioral, and genetic data from 1,570 healthy young adults (ages 18–35, mean age: 21.3 ± 2.73 years; females: 526 (58.6%)). All data were acquired on matched 3T Tim Trio scanners using 12-channel phased-array head coils. Resting-state fMRI used gradient-echo EPI sequence (TR = 3000 ms, TE = 30 ms, flip angle = 85°, 3.0 mm isotropic voxels, 47 interleaved axial slices aligned to AC-PC plane). Subjects were instructed to remain still, stay awake, and keep eyes open. 4 resting-state fMRI runs per subject were acquired, each with 120 valid whole-brain volumes (about 6 minutes 12 seconds per run). Age, sex, cognition, and education metrics were utilized for GSP-specific PI-FC+ model training.

### Healthy Brain Network (HBN)

We utilized data from 1,078 subjects from this Child Mind Institute initiative collecting data from 10,000 New York City area children and adolescents (ages 5–21, mean age: 10.5 ± 3.6 years; females: 687 (63.7%)) for transdiagnostic pediatric mental health research. Three types of fMRI data were incorporated: resting-state fMRI, movie-watching (naturalistic viewing) fMRI, and PEER calibration scans. Data was primarily acquired at Rutgers University Brain Imaging Center using 3T Siemens Tim Trio scanner with 32-channel head coil and CMRR simultaneous multi-slice EPI sequence (TR = 800 ms, TE = 30 ms, flip angle = 31°, 2.4 mm isotropic voxels, 60 slices, multi-band acceleration factor = 6). Resting-state data consisted of two 5-minute runs. Movie-watching fMRI included clips from ‘Despicable Me’ and ‘The Present’. Comprehensive phenotypic variables including demographics, cognitive assessments, and clinical/behavioral questionnaires were employed in HBN- specific PI-FC+ model training.

### Chinese Human Connectome Project (CHCP)

To incorporate Eastern population data and enhance cross-cultural comparisons, we included data from 304 healthy adult subjects (age range: 18–79 years, mean age: 34.0 ± 18.1 years; females: 163 (53.6%)), primarily Han Chinese individuals. Imaging data were acquired on 3T Siemens Prisma MRI scanner. Rs-fMRI was acquired using gradient-echo EPI sequence (TR = 710 ms, TE = 30 ms, flip angle = 52°, 2×2×2 mm^3^ voxels, 72 axial slices, acceleration factor = 8). Subjects typically underwent scanning over two days, resulting in 4 rs-fMRI runs per subject (totaling about 30 minutes). Behavioral and demographic measures including task-specific accuracies and reaction times assessing working memory, language processing, emotion recognition, relational reasoning, social cognition, and reward processing were utilized for CHCP-specific PI-FC+ model training.

### Clinical Validation Datasets

#### Human Connectome Project for Early Psychosis (HCP-EP)

To assess PI-FC+ framework performance in early psychosis, we utilized data from 134 subjects (90 patients, 44 healthy controls) from young adults (ages 16–35, mean age: 23.5 ± 4.0 years; females: 77 (62.6%)) within five years of psychosis onset, encompassing both affective and non-affective psychotic disorders. Data collection protocols were consistent with HCP-YA study protocols. Imaging data were acquired on 3T Siemens MAGNETOM Prisma scanners at Brigham and Women’s Hospital, McLean Hospital, and Indiana University using CCF 2016 template protocols. Rs-fMRI was acquired using gradient-echo EPI sequence (TR = 800 ms, TE = 37.00 ms, flip angle = 52°, 72 slices, FOV = 208 × 208 mm, 2mm isotropic resolution, multi-band acceleration factor = 8). 4 rs-fMRI runs were utilized per subject, each comprising 420 volumes. For cognitive inference analyses, the NIH Toolbox total cognition composite score (unadjusted) was utilized as the key cognitive measure. All data underwent HCP minimal processing pipeline.

#### Autism Brain Imaging Data Exchange (ABIDE)

We utilized data from 483 subjects (269 ASD, 214 typically developing controls, age range: 7–64 years, mean age: 16.3 ± 7.5 years; females: 108 (22.3%)) from seven contributing sites: Olin Neuropsychiatry Research Center, University of Michigan, Yale School of Medicine, Kennedy Krieger Institute, NYU Langone Medical Center, UCLA, and California Institute of Technology. Each subject underwent rs-fMRI and T1-weighted structural MRI scans with site-specific imaging parameters. Comprehensive phenotypic data including diagnostic information and intelligence measures were provided. For cognitive prediction analyses, Full Scale IQ (FIQ) derived from Wechsler Abbreviated Scale of Intelligence (WASI) was utilized, including only subjects with available FIQ scores.

### Strategic Research Program for the Promotion of Brain Science (SRPBS)

To evaluate our framework on a large-scale, clinically diverse Japanese cohort, we utilized data from 727 subjects (222 MDD patients, 495 healthy controls, age range: 18-80 years, mean age: 42.6 ± 14.4 years; females: 388 (54.1%)) selected based on data availability. Data were collected across eight research sites in Japan using 14 distinct 3T MRI scanners from three manufacturers (Siemens, Philips, General Electric). Rs-fMRI data were acquired using T2-weighted EPI sequences with harmonized protocols (TR = 2500 ms, TE = 30 ms, flip angle = 80°, voxel sizes = 3.3×3.3×3.2 mm with 0.8 mm gap, 40 slices, 10 minutes duration, 240 volumes). Subjects with mean framewise displacement ≥ 0.3 mm were excluded to standardize quality across 8 sites (multi-vendor)^86^.

### Generalized Anxiety Disorder (GAD)

To investigate neural correlates of generalized anxiety disorder and evaluate PI-FC+ framework clinical performance, we utilized a dataset from Henan Provincial People’s Hospital, Zhengzhou, China. The dataset comprised 87 subjects: 45 individuals diagnosed with GAD according to DSM-5 criteria and 42 age-matched healthy controls (age range: 18–59 years, mean age: 35.6 ± 10.8 years; females: 60 (69.0%)), with comprehensive clinical assessment and screening to exclude comorbid neurological conditions, substance abuse, and other major psychiatric disorders. Neuroimaging data were acquired using 3T Siemens Prisma MRI scanner with 64-channel head-neck coil. Structural T1-weighted images used 3D MPRAGE sequence (TR = 2530 ms, TE = 2.96 ms, TI = 1100 ms, flip angle = 7°, 1 mm isotropic voxels, parallel imaging factor = 2). Rs-fMRI used 2D EPI sequence (TR = 3000 ms, TE = 30 ms, flip angle = 85°, slice thickness = 3 mm, matrix = 72 × 72, 47 axial slices, anterior-posterior phase encoding).

### Data preprocessing

#### fMRI data preprocessing (fMRIprep)

For 6 datasets (Cam-CAN, GSP, HBN, CHCP, SRPBS, GAD), we performed preprocessing using fMRIPrep 23.2.1^87^. T1-weighted images underwent N4BiasFieldCorrection (ANTs 2.5.0), skull stripping with the Nipype implementation of antsBrainExtraction (OASIS30ANTs target), tissue segmentation with FSL FAST^88^, and nonlinear normalization to MNI152NLin2009cAsym via antsRegistration, with templates from TemplateFlow^89^. For each BOLD run, fMRIPrep generated a reference volume, estimated head motion using FSL MCFLIRT, and performed BOLD-to-T1w coregistration using FreeSurfer mri_coreg followed by FSL FLIRT with boundary-based registration (BBR)^90^. Susceptibility-distortion correction used fieldmaps when available or fieldmap-less SDC-SyN otherwise. Spatial transforms for motion, distortion, coregistration, and normalization were concatenated and applied in a single resampling step (cubic B-spline). Unless otherwise noted, we used volumetric outputs in MNI152NLin2009cAsym space. The ABIDE data were downloaded from the Amazon S3 repository as fMRIPrep-preprocessed files. Following fMRIPrep preprocessing, we applied systematic confound regression using nilearn’s load confounds function (nilearn.interfaces.fmriprep.load_confounds) with the following strategy: motion parameters (full 24-parameter model including 6 rigid-body parameters, their temporal derivatives, and squared terms), white matter (WM) and cerebrospinal fluid signals (CSF), and global signal. Additional nuisance regressors included a constant term, linear trend (t), and quadratic trend (t^2^) to account for scanner drift. No temporal band-pass filtering was applied. To avoid the impact of GSR on a short time scale, we generated two versions of the data: one with GSR and one without GSR. Traditional FC baselines used GSR.

#### fMRI data preprocessing (HCP series dataset)

For HCP-YA and HCP-D, we used outputs from the HCP minimal preprocessing pipelines^52^. Within that framework, resting-state and film fMRI underwent gradient-distortion, motion, and EPI-distortion correction; BOLD-to-T1w coregistration with BBR and single-subject spatial ICA followed by automated component classification with FIX using an HCP-trained classifier; components labeled as noise were removed via regression (sICA-FIX)^91,92^. Our analyses used the volumetric “MNINonLinear” outputs after sICA-FIX denoising. No additional nuisance regression (including 24-parameter motion, or WM/CSF signals) or censoring was applied to the sICA-FIX–denoised data. For HCP-EP and other task fMRI in HCP series dataset, data were processed with the HCP minimal preprocessing pipelines only. We applied the same nuisance-regression strategy as our fMRIPrep datasets.

### fMRI data preprocessing (UKBB dataset)

We used the centrally released preprocessed volumetric rs-fMRI data from UKBB, which underwent motion correction, grand-mean intensity normalization, EPI and gradient-distortion unwarping, and structured artifact removal via sICA-FIX^51,93^.

### Calcluation of functional connectome

Following our previous work^94^, we extracted time series from 271 whole-brain regions of interest (ROIs) and constructed 271 × 271 functional connectome matrices using pearson correlation. The 271 ROIs were derived from three complementary atlases to ensure comprehensive brain coverage: 200 ROIs from the Schaefer200 cortical parcellation^92,95,96^, 54 ROIs from the Melbourne Subcortex Atlas covering subcortical structures (3T version, scale IV)^97^, and 17 ROIs from the Buckner Atlas corresponding to cerebellar regions aligned with the 17 cortical networks^98^. All traditional FC analyses employed global signal regression as the comparison baseline, given its established effectiveness in reducing state-dependent confounds and enhancing individual-specific predictions^61,99^. This ensures that PI-FC improvements represent advances beyond this commonly used preprocessing approach rather than simple preprocessing effects. The FC extraction workflow was applied consistently across all datasets to ensure comparable FC representations for subsequent PI-FC framework training and validation.

### PI-FC framework

The PI-FC framework comprises three interconnected modules designed to extract stable, individual-specific FC representations while mitigating state-dependent fluctuations. The architecture includes: (i) a contrastive learning encoder for individual-specific feature extraction; (ii) a downstream prediction module for phenotype alignment; and (iii) a reconstruction module for stable FC pattern recovery (see Supplementary Fig. S5 for a schematic illustration).

**i. Contrastive learning module.** The core encoder employs a 1D convolutional neural network architecture to process functional connectome matrices. The network consists of five sequential layers with channel dimensions [271, 512, 256, 128, 64], utilizing 1D convolutions with ‘same’ padding, layer normalization, ELU activation functions, and 30% dropout regularization. Input FC matrices (36,585 upper-triangular elements) are first reconstructed into symmetric 271×271 matrices before processing. The encoder generates both shallow features (271×64 = 17,344 dimensions) and deep contrastive features (2,048 dimensions) through a fusion layer, resulting in concatenated representations of 19,392 dimensions.
**ii. Downstream prediction module.** A shared embedding layer (19,392 → 1,024) with layer normalization feeds task-specific heads for age regression and sex classification. For PI-FC+, additional dataset-specific behavioral heads (regression/classification) enable multi-phenotype learning.
**iii. Reconstruction module.** The reconstruction module aims to recover stable, individual-specific FC patterns from state-dependent inputs. Using the shared embedding features, it employs a deep feedforward network (19,392 → 5,120 → 36,585) with batch normalization, leaky ReLU activation, and hyperbolic tangent output activation to reconstruct the upper-triangular FC elements. This module enforces consistency by reconstructing both static long-term FC patterns and input short-term FC data.

### Multi-objective learning

The PI-FC framework employs a unified multi-task learning objective that simultaneously optimizes individual identification, phenotype prediction, and connectivity reconstruction:

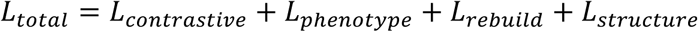

We implemented two variants of our framework: PI-FC (L_phenotype_ = L_age_ + L_sex_), which incorporates only age and sex prediction tasks in the downstream loss function during training, and PI-FC+ (L_phenotype_ = L_age_ + L_sex_ + L_behavior_), which encompasses all available behavioral phenotypes in the downstream prediction tasks to learn richer individual-specific representations.

### Individual-specific contrastive loss

The framework maximizes agreement between different temporal acquisitions from the same individual while minimizing cross-subject similarities using InfoNCE loss:

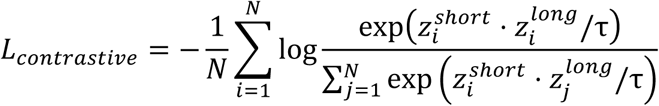

where 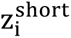 and 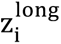 are 𝐿_2_-normalized embeddings from short- and long-duration FC data, and 𝜏 = 0.07 controls the temperature, with 𝑖 ∈ {1, … 𝑁} indexing the samples in the batch and 𝑗 ∈ {1, …, 𝑁} denoting the candidate indices (where 𝑗 = 𝑖 is the positive pair and 𝑗 ≠ 𝑖 are the negatives).

### Adaptive loss integration

To address optimization challenges across multiple diverse objectives, we implement a dynamic weighting strategy. During initial training (epochs 1-10), correlation-based losses dominate establishing stable feature relationships, followed by progressive integration of reconstruction objectives through a learnable parameter 𝜆:

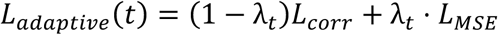

where λ_t_ increases from 0 to 0.5 over training epochs, balancing feature consistency and reconstruction fidelity.

### Structure-preserving constraint

To maintain inter-individual relationship patterns in the learned representations, we enforce structural consistency between feature-based and connectivity-based similarity matrices:

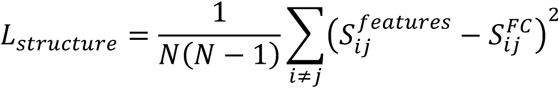

where S^features^ and S^FC^ represent cosine similarity matrices computed from learned embeddings and static FC patterns, respectively, with 𝑖, 𝑗 ∈ {1, …, 𝑁}, 𝑗 ≠ 𝑖 indexing the samples in the batch.

### Sample construction

Positive pairs comprise FC matrices from the same individual across different temporal windows (50-500 frames verus all frames) and task states. Negative pairs consist of FC matrices from different individuals across both resting and task states.

### Training details

Network parameters are initialized using Xavier uniform initialization, with bias terms set to zero. Training employs sequential stages: (1) contrastive pre-trainied, (2) joint multi-task optimization, and (3) dataset-specific fine-tuning over 500 epochs with early stopping (patience = 10 epochs). Batch sizes are set to 4,096 for most datasets and 2,048 for UK Biobank due to memory constraints. Adam optimization (learning rate = 1×10^-4^, weight decay = 1×10^-4^, ε = 1×10^-8^) with gradient clipping (max norm = 1.0) ensures stable training. All experiments are conducted on a single NVIDIA H800 GPU with 80GB memory. Dataset-specific normalization used pre-computed statistics enables robust cross-dataset generalization.

### Short-duration simulation and scan-time equivalence

We extracted sequential segments of 50 - 1,200 frames (TR = 0.72 s) from non-filtered, nuisance-regressed time series. At each duration, performance was computed over 100 independent outer-fold resamples. For scan-time equivalence, we linearly interpolated frames as a function of performance (SciPy interp1d, kind = ‘slinear’). Given FC duration x the minimum PI-FC (+) duration y achieving the same macro-averaged performance defined *y*(*x*); we reported y/x. Analyses were restricted to overlapping performance ranges to avoid extrapolation; 95% confidence intervals were obtained by paired bootstrap over seeds (B = 10,000); Confidence intervals (CIs) are reported in Supplementary Tables S1 but not displayed in the figure.

### Test-retest reliability

We assessed feature reliability using the two-way mixed, single-measure intra-class correlation ICC (3,1)^55,56^. For each connectome edge, ICC (3,1) was evaluated across the 4 runs. Under this model, ICC (3,1) quantifies the proportion of between-subject variance relative to total variance.

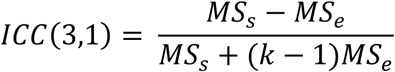

Where 𝑀𝑆_𝑠_ is the subjects mean square, 𝑀𝑆_𝑒_ is the residual mean square, and. 𝑘 is the number of runs (k = 4). For reference, the reliability of the run-averaged estimate corresponds to 𝐼𝐶(3, 𝑘) = (𝑀𝑆_𝑠_ −𝑀𝑆_𝑒_)/𝑀𝑆_𝑠_. Group summaries report mean ± SD across edges; uncertainty via percentile bootstrap over resamples (B = 10,000). Multiple comparisons across durations were controlled using Benjamini-Hochberg FDR at Q = 0.001.

### Arousal states

Each resting-state run from HCP-YA dataset was partitioned into an early period and a late period. The first and last 10 frames of the entire run were discarded to avoid steady state and end of scan artifacts. Consistent with prior reports^20,57^ of vigilance decline during prolonged resting-state acquisitions, the early and late segments were treated as proxies for relatively high and low arousal, respectively. FC matrices were then computed separately for each segment using both traditional FC and PI-FC. All downstream phenotypic predictions —feature normalization, model choice, hyperparameter search, cross validation strategy, and evaluation metrics— were applied identically to the early and late period.

### Dynamic brain states

To investigate how brain state variations influence functional connectome patterns, we performed dynamic functional connectome (dFC) analysis on resting-state fMRI data from the HCP-YA dataset. We applied a sliding window approach with 50-frame windows (36 seconds at TR = 0.72 s) and non-overlapping steps to extract time-resolved FC matrices from whole-brain ROI time series. Global signal regression was applied to enhance state differentiation. The resulting high-dimensional dFC data were reduced to 1,000 features using PCA to facilitate clustering analysis. K-means clustering was performed exclusively on the HCP-YA training set, systematically evaluating cluster numbers from 2 to 10, with the optimal number (k = 4) determined using the elbow method based on within-cluster sum of squares. This identified 4 predominant brain states with distinct occurrence frequencies (state 1: 40.0%, state 2: 28.6%, state 3: 16.4%, and state 4: 15.0%). For each identified brain state, we constructed state-specific FC matrices by averaging connectivity patterns across all time windows assigned to that particular state within each individual, yielding 4 state-specific FC matrices per subject representing characteristic connectivity patterns during each meta-state. To establish individual-specific reference patterns, we computed “long FC” for each subject by averaging connectivity across all available scanning sessions and paradigms (including all 4 resting-state runs and task-based fMRI sessions when available), representing their stable, trait-like functional architecture. We characterized population-level state structures by computing group-averaged connectivity matrices and quantified individual state preservation through Pearson’s r between each subject’s state-specific FC matrices and their corresponding long FC reference. Cross-state similarity structure preservation was evaluated by computing behavioral phenotype similarity matrices using Euclidean distances across 39 HCP-YA phenotypic measures and comparing their correspondence with connectivity-based similarity matrices derived from state-specific FC patterns using Frobenius norm correlations for both traditional FC and rebuild PI-FC patterns. The trained clustering model was subsequently applied to predict meta states in independent test set data for downstream validation analyses. Cross-state generalization followed two designs: train-on-one-state test-on-others and train-on-others test-on-held-out state, using the same nested-CV and models as other analyses; differences were assessed by paired t-tests across resamples.

### Cross-paradigm validation

To evaluate the cross-paradigm generalizability between traditional FC and PI-FC, we designed two validation strategies. In the first approach, models were trained on resting-state data and tested directly on task data without any retraining, simulating a realistic cross-paradigm transfer scenario. In the second approach, we created mixed-paradigm datasets by randomly selecting either resting-state or task data for both training and testing sets. Each subject contributed only one kind of data (resting-state or task) to ensure that no subject appeared in both training and testing sets.

### Phenotypes

To train, validate, and evaluate the PI-FC+ framework, we employed a hierarchical phenotyping strategy across different model configurations and datasets.

#### Pre-trained model 1

We performed deep behavioral characterization using HCP-YA dataset with 39 measures including three core domains: cognitive function (such as fluid intelligence, crystallized intelligence, working memory), personality (NEO Five-Factor Inventory), and mental health (such as sleep quality, substance use, mood assessments, life satisfaction, social relationships). A detailed list of phenotypes is provided in Supplementary Table S8.

#### Pre-trained model 2

For multi-dataset integration, we utilized harmonized phenotypes across diverse populations including cognitive function (such as fluid intelligence, working memory, processing speed, executive control), personality (neuroticism, behavioral inhibition/activation), and mental health (such as depression, anxiety, sleep, substance use). Dataset-specific measures totaled 124 variables across HCP-D (56), UKBB (25), GSP (9), HBN (22), CHCP (22) and CAM-CAN (2). A detailed list of phenotypes is provided in Supplementary Table S8.

### Clinical validation

We validated clinical utility across patient cohorts spanning early psychosis (HCP-EP), autism spectrum disorder (ABIDE), depression (SRPBS), and generalized anxiety disorder (GAD), examining diagnostic classification and cognitive measures including IQ scores and executive function assessments. All phenotypic variables are detailed in Supplementary Table S8.

### Fingerprinting

Fingerprinting analyses^14^ assessed the preservation of person-specific connectivity signatures across tasks and temporal scales using 200 independent HCP-YA subjects (random selected and repeat 100 times). We evaluated identification accuracy within-task (across runs of identical paradigms) and cross-task (between different experimental conditions) using resting-state and seven task paradigms (gambling, motor, working memory, emotion, language, relational, social) with 100-frame segments (72 s). For each subject, Pearson’s r were computed between connectivity patterns in source and target conditions, with identification succeeding when the highest correlation correctly matched the same individual. Traditional FC (36,585 upper-triangular matrix elements) was compared against contrastive PI-FC individual embeddings (2,048 dimensions) following z-score normalization. To facilitate visualization, we applied UMAP to obtain a low-dimensional embedding (n_neighbors = 15, min_dist = 0.1, cosine metric) for subjects across eight task conditions, and quantified separation with silhouette coefficients for identity versus task. UMAP preserves local neighborhood structure by constructing a k nearest-neighbor fuzzy graph and optimizing a low-dimensional layout to match those neighborhood relations^64^. Identification was repeated across multiple random seeds with different subject selections, and performance differences between traditional FC and PI-FC were assessed using paired t-tests with all analyses employing fixed random seeds for reproducibility.

### Downstream prediction pipeline

The PI-FC framework enables three types of downstream prediction tasks: behavioral phenotype prediction, psychiatric disorder classification, and direct phenotype inference across independent datasets.

### Machine learning model (input features)

We used two feature families: (i) traditional FC, computed as described in Functional connectome extraction; and (ii) PI-FC embeddings learned from the same FC inputs. In PI-FC, the encoder yields shallow features (271×64 = 17,344) and deep contrastive features (2,048), fused to 19,392 dims; concatenating the 1,024-dim shared embedding from the downstream prediction module gives a 20,416-dim representation. If a subject had multiple runs, we computed features per run with the same steps and then averaged them in clinical application, but in HCP-YA dataset, the experiment required keeping them separate (such as early vs. late, dynamic states). All features were z-scored within each training fold, with the same scaler applied to the test fold to avoid data leakage.To improve prediction performace, we used the GSR version of the FC features. This choice is consistent with previous large-scale studies showing that GSR enhances inter-individual variability in functional connectomes and improves the accuracy of behavioral predictions derived from resting-state fMRI^61,99^.

### Machine learning model (kernel ridge regression)

For each non-imaging continuous phenotype, we trained a kernel ridge regression (KRR) model using nested cross-validation on the full dataset. Each random split was partitioned into 5 outer folds, with family members (where applicable, e.g., in the HCP-YA dataset) kept within the same fold to avoid relatedness leakage. Within each outer-train fold we performed 5-fold inner CV over a logarithmic grid of the 𝐿_2_ regularization hyperparameter 𝜆. To choose 𝜆, validation performance was evaluated using the Pearson’s r, and the selected 𝜆 was then used to refit KRR on the entire outer-train fold and evaluated on the held-out outer-test fold. Across repetitions, test performance was summarized with Pearson’s r.

### Machine learning model (logistic regression)

For binary outcomes (clinical diagnosis, sex), we used 𝐿_2_regularized logistic regression under the same 5 × 5 nested CV scheme. Each random split was partitioned into 5 outer folds, with family members (where applicable, e.g., in the HCP-YA dataset) kept within the same fold to avoid relatedness leakage. In the inner loop, the inverse penalty C was tuned over a logarithmic grid, with model selection based on bACC on the validation folds. The optimal C was then usded to refit the model on the entire outer-train fold and evaluated on the outer-test fold. Test performance was reported as bACC and AUC.

### Cross-validation strategies

Model performance was evaluated using a rigorous 100-iteration, 5-fold nested cross-validation approach to ensure robust and stable predictions. This extensive validation strategy was consistently applied across all phenotype prediction tasks, including the behavioral measures from HCP-YA datasets. For multi-site datasets (ABIDE, SRPBS), leave-one-site-out cross-validation (LOSO CV) was additionally employed to assess generalizability across acquisition sites and scanner heterogeneity. For single-site dataset (HCP-EP, GAD), we applied the standard 5-fold nested cross-validation. No ComBat harmonization was applied to avoid leakage; robustness was assessed via LOSO and stacking.

### Multi-source ensemble strategy

For clinical applications, we implemented a stacking ensemble approach using PI-FC+ (*pre-trained model 2*, *stacking*). For each target clinical cohort, all available PI- FC+ models pre-trained on datasets with no subject overlap were identified. Individual model predictions were then averaged to generate final ensemble predictions, leveraging complementary strengths learned from diverse populations and acquisition protocols.

### Direct inference capability

The pre-trained PI-FC+ framework enables direct phenotype inference in completely independent clinical centers without requiring additional training data. This zero-shot inference capability was validated for age, sex, and cognitive ability across multiple clinical cohorts that were entirely separate from pre-trained datasets. We performed inference using the cognitive scales listed in Supplementary Table S9.

### Covariate control

For behavioral phenotypes prediction using KRR, we residualized targets for age and sex within each training fold and applied the fitted nuisance model to the corresponding test targets (no leakage). For clinical classification tasks, bACC and AUC served as primary performance metrics, while Pearson’s r was used for continuous phenotype prediction tasks.

## Data Availability

Most datasets in our study are available under their respective data-use terms. Raw and processed 3T and 7T fMRI from the Human Connectome Project (HCP Young Adult S1200) are available at https://www.humanconnectome.org/study/hcp-young-adult/document/1200-subjects-data-release. UK Biobank imaging (including brain MRI) is available to researchers upon application to UK Biobank (https://www.ukbiobank.ac.uk/about-our-data/types-of-data/imaging-data/). Cambridge Centre for Ageing and Neuroscience (Cam-CAN) data can be requested via the Cam-CAN Data Repository (https://cam-can.mrc-cbu.cam.ac.uk/dataset/). Brain Genomics Superstruct Project (GSP) open-access data are available from Harvard (Buckner Lab / Neuroinformatics Research Group) (https://bucknerlab.fas.harvard.edu/gsp). Healthy Brain Network (HBN) MRI datasets are available via the HBN data portal / NITRC (https://data.healthybrainnetwork.org/main.php). HCP Lifespan Development (HCP-D) data are distributed through the NIMH Data Archive (NDA) (https://www.humanconnectome.org/study/hcp-lifespan-development/data-releases). Chinese Human Connectome Project (CHCP) information and data are provided by the project (https://www.scidb.cn/en/detail?dataSetId=f512d085f3d3452a9b14689e9997ca94). Additional evaluation cohorts include ABIDE (Autism Brain Imaging Data Exchange) and HCP Early Psychosis (HCP-EP; distributed via the NIMH Data Archive), and SRPBS multi-disorder MRI datasets (available from the SRPBS consortium under controlled-access terms). Other raw data can be requested from the corresponding author.

## Code Availability

Code for PI-FC/PI-FC+ training, evaluation, and downstream could be found at a public repository (https://github.com/hlpureboy/PI-FC). The implementation is based on Pytorch with supporting libraries including torchvision, numpy, scipy, pandas, scikit-learn, lightly, nilearn, and h5py. Detailed installation instructions and environment files are provided in the repository.

## Acknowledgement

This research was conducted using data from multiple open-access neuroimaging initiatives. Data were provided in part by the Human Connectome Project, WU-Minn Consortium (PIs: David Van Essen and Kamil Ugurbil), and the Lifespan Human Connectome Project Development (PIs: Essa Yacoub and David Van Essen). Additional data were obtained from the UK Biobank under application no. 85139 (PI: Rory Collins), the Healthy Brain Network (see http://www.healthybrainnetwork.org for full donor list), and the Human Connectome Project for Early Psychosis, supported by the National Institute of Mental Health (Award U01MH109977; Release 1.1, DOI: 10.15154/1522899). We further acknowledge the Cambridge Centre for Ageing and Neuroscienc, supported by the Biotechnology and Biological Sciences Research Council, and the Brain Genomics Superstruct Project (Harvard University and Massachusetts General Hospital; PIs: Randy Buckner, Joshua Roffman, and Jordan Smoller), with support from the Center for Brain Science Neuroinformatics Research Group, the Athinoula A. Martinos Center for Biomedical Imaging, and the Center for Human Genetic Research. Data were also provided by the Chinese Human Connectome Project (PI: Jia-Hong Gao), funded by the Beijing Municipal Science & Technology Commission, the Chinese Institute for Brain Research (Beijing), the National Natural Science Foundation of China, and the Ministry of Science and Technology of China; and the DecNef Project Brain Data Repository (https://bicr-resource.atr.jp/srpbsopen/), conducted as part of the Japanese Strategic Research Program for the Promotion of Brain Science supported by AMED. Finally, data from the Autism Brain Imaging Data Exchange, generated as part of the PsychENCODE Consortium (see https://doi.org/10.7303/syn24240356 for full list of grants and PIs), were obtained through the Autism BrainNet (Simons Foundation) and the NIH NeuroBiobank (University of Maryland Brain and Tissue Bank). This work was supported by the National Key R&D Program of China (2023YFC2414200), STI2030-Major Projects (2022ZD0211900), Beijing Nova Program (20250484761), the Natural Science Foundation of China (82171543, 8244102, 8244102, 82371934).

## Author contributions

A.L., Y.Y. and M.W. led the project. A.L. and Y.P. conceived the concept and designed the experiments. Y.P. developed the PI-FC computational implementation. Y.P. and X.T. conducted the analyses, with support from Y.H., S.H., T.Z., J.X., T.G., and Q.W. Data preprocessing was completed by Y.P., X.T., C.D., X.L., K.L., and Y.G., X.Z., L.W., Y.T., and B.L. provided feedback on the analyses. M.W. contributed clinical fMRI data. Y.Y. provided critical suggestions for the work. Y.P. and A.L. created the figures and drafted the manuscript.

